# Nanopore quality score resolution can be reduced with little effect on downstream analysis

**DOI:** 10.1101/2022.03.03.482048

**Authors:** Martín Rivara-Espasandín, Lucía Balestrazzi, Guillermo Dufort y Álvarez, Idoia Ochoa, Gadiel Seroussi, Pablo Smircich, José Sotelo-Silveira, Álvaro Martín

## Abstract

We investigate the effect of quality score information loss on downstream analysis from nanopore sequencing FASTQ files. We polished denovo assemblies for a mock microbial community and a human genome, and we called variants on a human genome. We repeated these experiments using various pipelines, under various coverage level scenarios, and various quality score quantizers. In all cases we found that the quantization of quality scores cause little difference on (or even improves) the results obtained with the original (non-quantized) data. This suggests that the precision that is currently used for nanopore quality scores is unnecessarily high, and motivates the use of lossy compression algorithms for this kind of data. Moreover, we show that even a non-specialized compressor, like gzip, yields large storage space savings after quantization of quality scores.

## 1 Introduction

The precision for quality score representation in FASTQ files has historically been relatively large, where each score takes a value within a set of more than 40 possible values. The precise definition of this set depends on the sequencing technology; sequencers from Oxford Nanopore Technologies^1^ use a set of 94 different values (represented in FASTQ files as ASCII codes 33 to 126). This high precision makes quality score sequences hard to compress and, thus, most of the information stored in a losslessly compressed FASTQ files corresponds to quality scores (substantially more than the genomic information itself). With growing concerns about the economics of storage space and bandwidth for transmission of genomic data, this policy is being questioned. Illumina, for example, has reduced the set of possible values for quality scores on their most modern equipment to just 4 different values [1] and suggests similarly quantizing the values produced by other Illumina devices [2]. In the context of compression algorithms for short-read sequencing data, it has also been observed that it is possible to use lossy compression algorithms, in which the decompressed data may differ to some extent from the original, without generally affecting the results of commonly used pipelines [3, 4, 5].

Nanopore sequencing, however, is a much more recent technology and few specific data compressors suitable for nanopore data are available, developed by our group [6, 7] and others [8, 9]. Moreover, the lossy compression of quality scores for nanopore data has only been explored in [9], where the impact of quality score information loss is assessed for some downstream analyses. Specifically, in [9], it is shown that this information loss has non or little impact on the construction of consensus sequences with Racon [10] for long CHM13 reads, either for HiFi or for Nanopore data. The impact is also shown to be small for variant calling accuracy with HiFi reads, evaluated with DeepVariant [11] on HG002 HiFi reads against the human reference genome GRCh38; variant calling for nanopore data is not evaluated.

In this paper we evaluate the impact of nanopore quality score quantization on various downstream applications, by comparing the results obtained with certain non-quantized nanopore data sets, referred to as *original data*, and with quantized versions of the same data sets. Particularly we selected data sets relevant for microbiology and human genetics, both areas of high interest and increasing use of this sequencing technology. The quantizers that we evaluate are mostly simple functions, where the set of possible quality scores is partitioned into a small set of disjoint intervals, each of which is mapped to a fixed value. For example, one such a quantizer maps all quality scores between 0 and 7 to the fixed score 5, and maps all quality scores larger than 7 to the fixed score 15. We also evaluate a slightly more complex scheme, which uses higher quantization resolution for quality scores associated to repetitive regions of a read. A precise definition of all evaluated quantizers is given in Section 2.1.

To asses the effect of quality score quantization on typical bioinformatic applications, we carried on assembly polishing for the ZymoBIOMICS Microbial Community Standard (Zymo Research Corporation, Irvine, CA, USA. Product D6300, Lot ZRC190633) nanopore reads using Racon [10] both alone and combined with Medaka,^2^ and MarginPolish [12] both alone and combined with HELEN [12]. We also evaluated the quality of human genome assembly from nanopore reads polished with MarginPolish both alone and combined with HELEN, for several coverage levels. For variant calling we executed PEPPER-Margin-DeepVariant [13] on sample HG003 for several coverage level scenarios. Details on the methods for evaluating assembly polishing and variant calling pipelines are presented in sections 2.2 and 2.3, respectively.

As shown in Section 3, all the experimented pipelines yield very similar performance on the original and quantized versions of each data set. These results complement and reinforce the aforementioned conclusion in [9], indicating that the effect of information loss caused by quality score quantization is not significant in practice. For the assembly polishing of a mock microbial community, setting all quality scores to the fixed value 10 results in a number of mismatches that is, on average over three independent runs, less than 1.2 % higher that that obtained with the original data. A quantizer that uses 4 different values for quality scores results in a number of mismatches that is, on average, even smaller than that obtained with the non-quantized data. This same quantizer, applied on human data, results in a number of mismatches that is essentially equivalent to that obtained with the original data; even for a very low coverage (10 % of the original reads), the increment in the number of mismatches is about 0.7%. Moreover, we show that the quantization of quality scores yields very large improvements in compression performance even for traditional non-specialized compressors such as gzip (see sections 2.4 and 3). For example, we report on human variant calling results at 90× coverage where using 8 different values for quality scores yields precision and recall metrics for single nucleotide polymorphisms that differ in less than 10^−4^ from those obtained with the original (non-quantized) data set, while saving more than 33% of the required storage space. Even using only 2 different values for quality scores, which saves almost 70% of storage space, the precision differs in less than 10^−4^ and the recall in less than 10^−3^ from those obtained with the original data set.

## 2 Methods

### 2.1 Quantization of quality scores

We tested various quantizers, each mapping quality scores to values on a (small) set referred to as the *quantization alphabet*. We denote by *Q*_*i*_ a quantizer for a quantization alphabet of size *i, i* > 1, where the quantization *Q*_*i*_(*x*) of a quality score *x* depends solely on *x*. The specific definitions of *Q*_2_, *Q*_4_, and *Q*_8_ are presented in tables 1, 2, and 3, respectively. All these quantizers collapse a large set of high quality scores into a single value, and define a finer partition for lower scores, which occur more frequently [14]. We also tested constant quantizers, which map every quality score to a fixed prescribed value. We denote by *F*_*z*_ a quantizer that assigns the value *F*_*z*_(*x*) = *z* to every quality score *x*. In our experiments we take *z* = 10, which is a common threshold for filtering low quality reads (see, e.g., [13]).

**Table 1:** Definition of quantizer *Q*_2_.

| Quality scores | Quantized quality score |
| --- | --- |
| 0...7 | 5 |
| 8...93 | 15 |

**Table 2:** Definition of quantizer *Q*_4_.

| Quality scores | Quantized quality score |
| --- | --- |
| 0...7 | 5 |
| 8...13 | 12 |
| 14...19 | 18 |
| 20...93 | 24 |

**Table 3:**
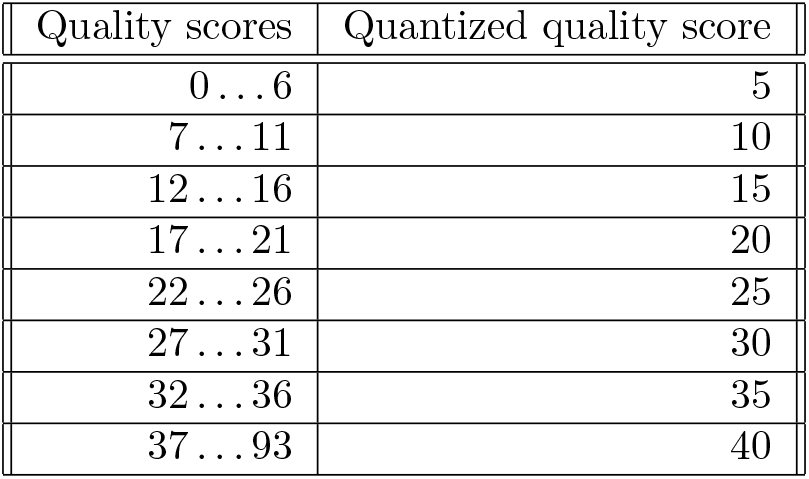
Definition of quantizer *Q*_8_.

| Quality scores | Quantized quality score |
| --- | --- |
| 0...6 | 5 |
| 7...11 | 10 |
| 12...16 | 15 |
| 17...21 | 20 |
| 22...26 | 25 |
| 27...31 | 30 |
| 32...36 | 35 |
| 37...93 | 40 |

In addition, since repetitive patterns of bases are particularly difficult to sequence by nanopore technologies [14], we evaluate quantizers where the quantization of a quality score *x* in a position *i* of a read depends not only on *x* but also on the bases called in positions close to *i*. We say that a substring of the base call sequence of a read is a *repetitive sequence* if it is comprised of either *h* consecutive copies of the same symbol, i.e., an homopolymer of length *h*, or *d* consecutive copies of the same pair of symbols, i.e., a repeat of *d* dinucleotides. For two quantization functions, *f*, *f*′, we denote by ⟨*f, f′*⟩_*δ*_ a quantizer that quantizes *x* by *f* (*x*), if the position *i* of the quality score *x* in a read is more than *δ* bases away from a repetitive sequence in this read, and by *f′*(*x*) otherwise. For example, the quantizer ⟨*F*_10_, *Q*_8_⟩_*δ*_ sets all quality scores away from repetitive regions to the fixed value 10, and applies *Q*_8_ to quality scores that are within or close to repetitive regions. For the experiments reported in Section 3, the parameters *h*, *d*, and *δ* are set to 5, 4, and 5, respectively.

### 2.2 Assembly polishing

We evaluated the impact of quality score quantization on the genome assembly polishing for the Zymo-BIOMICS Microbial Community Standard (Zymo Research Corporation, Irvine, CA, USA. Product D6300, Lot ZRC190633). For the experiments we used the data generated by the authors of [15] on Oxford Nanopore GridION, R10.3 pore^3^. The organisms in this community are 8 bacteria, each present at 12%, and 2 yeasts, each present at 2%. We assembled the genomes with Flye [16] and kept the 8 largest contigs, which correspond to the 8 bacteria. We polished this raw assembly using various polishing pipelines:

- *Racon*: 2 iterations of Racon [10].
- *Medaka*: 2 iterations of Racon followed by Medaka.^4^
- *MP*: An execution of MarginPolish [12].
- *Helen*: An execution of MarginPolish followed by HELEN [12].

Both Racon and MarginPolish are polishers that make use of quality scores. Medaka and HELEN, on the other side, are designed to refine the base polish obtained by Racon and MarginPolish, respectively.

We tested each polishing pipeline on the original (non-quantized) data, and quantized versions using *F*_10_, *Q*_2_, *Q*_4_, and *Q*_8_. To account for randomness within the executed algorithms we ran three executions of each combination of polisher and data set.

As reference for assessment of the results we used a combination of scaffolds obtained by SPAdes [17] from Illumina reads as described in [15].^5^ We evaluated the quality of the final assembly for each run of a polishing pipeline on a data set using MetaQUAST [18] to obtain the number of mismatches per kilobase pairs (kbp) with respect to the combined reference.

To analyze the results of assembly polishing on quantized quality scores data in a different setting, we followed some of the experiments reported in [12] for human genome assembly. Specifically, we polished the assembly generated by wtdbg2 [19] for one of the flows for sample HG00733 [20] with the polishing pipelines MP and Helen. These polishing pipelines were executed both for the orginal FASTQ files and for the same data quantized with *Q*_4_, and we compared the number of mismatches per 100 kbp with respect to the reference GRCh38 using QUAST [21]. We carried on this comparison for several coverage scenarios, which we obtained by randomly selecting a fraction (10 %, 20 %, 60 %, and 100 %) of the dataset reads. We repeated each execution three times to account for algorithms randomization.

### 2.3 Variant calling

We compared the nanopore variant calling performance of PEPPER-Margin-DeepVariant on sample HG003, as reported in [13], against variant calling on quantized versions of the same data. We performed this comparison at various coverages, ranging from 20× to 90×, and for various quantizers described in Section 2.1.

Following [13], we used GIAB v4.2.1 truth set against GRCh38 reference, and GIAB stratification v2.0 files with hap.py^6^ to derive stratified variant calling results. As a result, for each coverage and each quantized version of the data, we obtain a classification of the variants reported by PEPPER-Margin-DeepVariant into two categories: true-positives (TP), and false-positives (FP), and a set of true variants that PEPPER-Margin-DeepVariant failed to identify, i.e., false-negatives (FN). These variants are reported separately for single nucleotide polymorphisms (SNPs) and insertions / deletions (INDELs). From these reports we calculate three statistics that summarize the performance of PEPPER-Margin-DeepVariant for each specific type of variant (SNP/INDEL), on various coverage and quantization settings:

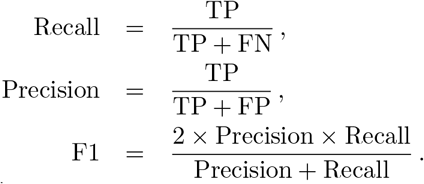

### 2.4 Compressibility

To evaluate the improvement in compressibility obtained by quantizing the quality scores of nanopore FASTQ files, we used sample HG003 (the same data set used for variant calling evaluation), which consists of three FASTQ files that add up to approximately 520 GB. We compressed the original data set and quantized versions of this data set using the general purpose compressor gzip^7^. For each evaluated quantizer we calculated the *compression ratio*, defined as the quotient between the size of the compressed data set and the size of the original data set (smaller ratios correspond to better compression performance).

## 3 Results

Table 4 presents the number of mismatches per 100 kbp for the assembly of a mock microbial community with respect to a combined reference (see Section 2.2), for various combinations of assembly polishing algorithm and quality score quantizer. The number of mismatches, averaged over three independent runs, is very similar among every combination of polisher and quantizer, with differences that are generally smaller than those among different runs of the same polisher and quantizer, which arise from algorithm randomization in the polishing pipeline. Comparing the average number of mismatches obtained with a certain polisher using quantized data and using the original data, we observe that the maximum difference is obtained for Medaka with the most heavy quantizer, *F*_10_. Even in this case, the difference represents only 1.2% of the average number of mismatches obtained with the original data. Notably, the results for the quantizer *Q*_4_ are even better than those obtained with the original data for all the evaluated polishing pipelines.

**Table 4:**
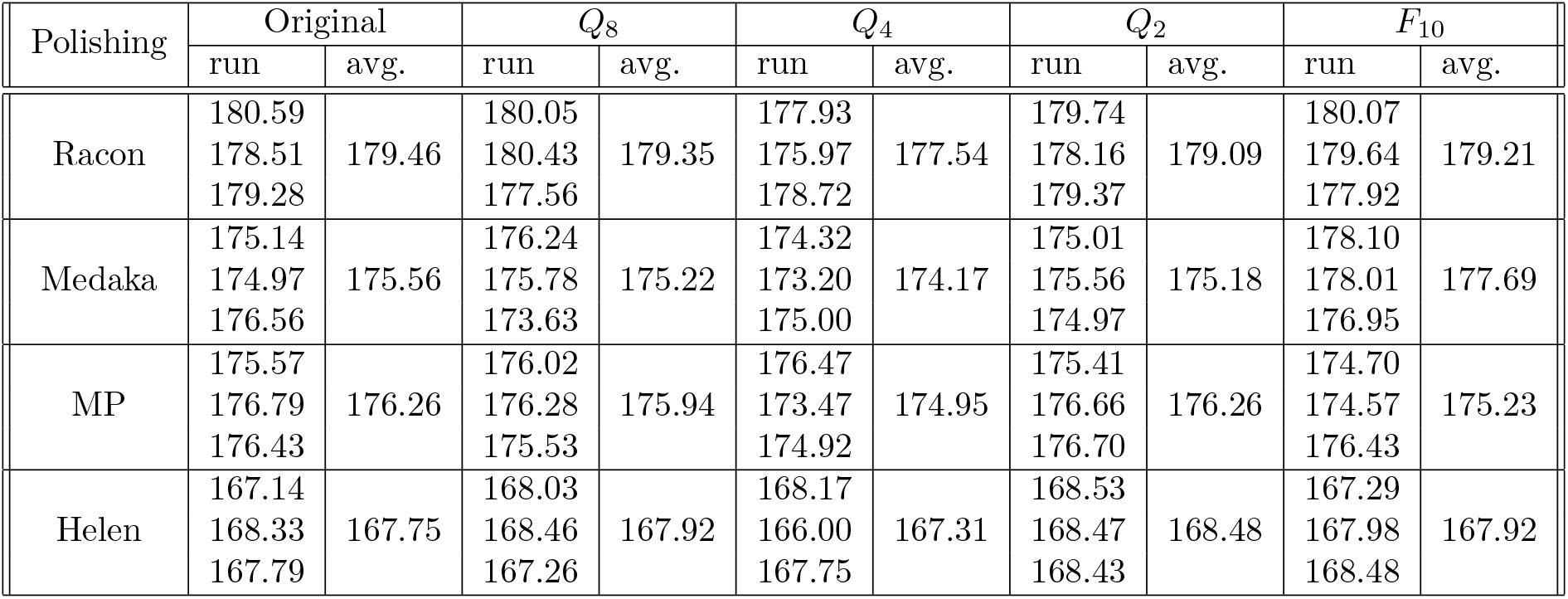
Number of mismatches per 100 kbp for the assembly of a mock microbial community, with respect to a combined reference, for various combinations of assembly polishing algorithm and quality score quantizer. For each polishing algorithm and each quantizer we show the result for three independent runs and the average of these three results.

| Polishing | Original | | $Q_8$ | | $Q_4$ | | $Q_2$ | | $F_{10}$ | |
| --- | --- | --- | --- | --- | --- | --- | --- | --- | --- | --- |
|  | run | avg. | run | avg. | run | avg. | run | avg. | run | avg. |
| Racon | 180.59 | 179.46 | 180.05 | 179.35 | 177.93 | 177.54 | 179.74 | 179.09 | 180.07 | 179.21 |
|  | 178.51 |  | 180.43 |  | 175.97 |  | 178.16 |  | 179.64 |  |
|  | 179.28 |  | 177.56 |  | 178.72 |  | 179.37 |  | 177.92 |  |
| Medaka | 175.14 | 175.56 | 176.24 | 175.22 | 174.32 | 174.17 | 175.01 | 175.18 | 178.10 | 177.69 |
|  | 174.97 |  | 175.78 |  | 173.20 |  | 175.56 |  | 178.01 |  |
|  | 176.56 |  | 173.63 |  | 175.00 |  | 174.97 |  | 176.95 |  |
| MP | 175.57 | 176.26 | 176.02 | 175.94 | 176.47 | 174.95 | 175.41 | 176.26 | 174.70 | 175.23 |
|  | 176.79 |  | 176.28 |  | 173.47 |  | 176.66 |  | 174.57 |  |
|  | 176.43 |  | 175.53 |  | 174.92 |  | 176.70 |  | 176.43 |  |
| Helen | 167.14 | 167.75 | 168.03 | 167.92 | 168.17 | 167.31 | 168.53 | 168.48 | 167.29 | 167.92 |
|  | 168.33 |  | 168.46 |  | 166.00 |  | 168.47 |  | 167.98 |  |
|  | 167.79 |  | 167.26 |  | 167.75 |  | 168.43 |  | 168.48 |  |

Table 5 compares the number of mismatches per 100 kbp for a human genome assembly obtained with the original (non-quantized) data and the data obtained by applying *Q*_4_ to quality scores. The comparison is performed for the polishers MP and Helen for the full data set (100 %), and for downsampled versions of the data consisting of 10 %, 20 %, and 60 % of the reads in the full data set (see Section 2.2). For a 20 % or larger fraction of the data, the results obtained after quantization are essentially equivalent to those obtained with the original data. Even for a very small fraction of the data (10 %), the percent increment in the number of mismatches incurred by quantization, averaged over three independent runs, is 0.01 % for MP and 0.7 % for Helen.

**Table 5:** Number of mismatches per 100 kbp obtained by MP and Helen for a human genome assembly, under various coverage scenarios, for the original (non-quantized) data and for the data obtained by applying *Q*_4_ to quality scores. For each combination of polisher, quantizer, and coverage, we show the result for three independent runs and the average of these three results.

| Polishing | Quantization | 10 % |  | 20 % |  | 60 % |  | 100 % |  |
| --- | --- | --- | --- | --- | --- | --- | --- | --- | --- |
|  |  | run | avg. | run | avg. | run | avg. | run | avg. |
| MP | Original | 1480.01 | 1480.01 | 935.24 | 935.24 | 132.89 | 132.89 | 100.32 | 100.32 |
|  |  | 1480.01 |  | 935.24 |  | 132.89 |  | 100.32 |  |
|  |  | 1480.01 |  | 935.24 |  | 132.89 |  | 100.33 |  |
| | $Q_4$ | 1480.01 | 1480.16 | 935.24 | 935.24 | 131.01 | 132.26 | 100.33 | 100.32 |
|  |  | 1480.24 |  | 935.24 |  | 132.89 |  | 100.32 |  |
|  |  | 1480.24 |  | 935.24 |  | 132.89 |  | 100.32 |  |
| Helen | Original | 1470.56 | 1460.03 | 907.05 | 907.05 | 125.94 | 125.94 | 99.65 | 99.65 |
|  |  | 1454.77 |  | 907.05 |  | 125.94 |  | 99.65 |  |
|  |  | 1454.77 |  | 907.05 |  | 125.94 |  | 99.65 |  |
| | $Q_4$ | 1470.56 | 1470.56 | 907.05 | 907.05 | 125.98 | 125.98 | 99.65 | 99.65 |
|  |  | 1470.56 |  | 907.05 |  | 125.98 |  | 99.65 |  |
|  |  | 1470.56 |  | 907.05 |  | 125.98 |  | 99.65 |  |

Tables 6 and 7 show the count of true-positives, false-positives, and false-negatives for SNPs and INDELs, respectively, obtained with PEPPER-Margin-DeepVariant for various quantization schemes and coverage levels. The tables also show the metrics Recall, Precision, and F1 calculated from these counts. Figures 1 and 2 show the complement to 1 of the F1 score, i.e., the difference 1 – F1, for SNPs and INDELs, respectively, for the same quantization schemes and coverage levels.

**Table 6:** Metrics TP, FP, FN, Recall, Precision, and F1 for the SNPs obtained with PEPPER-Margin-DeepVariant under various coverage scenarios, for the original (non-quantized) data and for the data obtained by applying various quality score quantizers.

| Coverage | Quantization | TP | FP | FN | Recall | Precision | F1 |
| --- | --- | --- | --- | --- | --- | --- | --- |
| 90× | Original | 3317116 | 10041 | 10376 | 0.9969 | 0.9970 | 0.9969 |
| | $Q_8$ | 3317064 | 10071 | 10428 | 0.9969 | 0.9970 | 0.9969 |
| | $Q_4$ | 3316599 | 9994 | 10893 | 0.9967 | 0.9970 | 0.9969 |
| | $Q_2$ | 3315132 | 10023 | 12360 | 0.9963 | 0.9970 | 0.9966 |
| | $F_{10}$ | 3310775 | 9522 | 16717 | 0.9950 | 0.9971 | 0.9961 |
| | $\langle Q_2, Q_8 \rangle_\delta$ | 3315867 | 10121 | 11625 | 0.9965 | 0.9970 | 0.9967 |
| 50× | Original | 3314614 | 13842 | 12878 | 0.9961 | 0.9958 | 0.9960 |
| | $Q_8$ | 3314564 | 14160 | 12928 | 0.9961 | 0.9957 | 0.9959 |
| | $Q_4$ | 3314059 | 14488 | 13433 | 0.9960 | 0.9956 | 0.9958 |
| | $Q_2$ | 3312857 | 14693 | 14635 | 0.9956 | 0.9956 | 0.9956 |
| | $F_{10}$ | 3310166 | 14337 | 17326 | 0.9948 | 0.9957 | 0.9952 |
| | $\langle Q_2, Q_8 \rangle_\delta$ | 3313530 | 14780 | 13962 | 0.9958 | 0.9956 | 0.9957 |
| 40× | Original | 3312510 | 20159 | 14982 | 0.9955 | 0.9940 | 0.9947 |
| | $Q_8$ | 3312554 | 20778 | 14938 | 0.9955 | 0.9938 | 0.9946 |
| | $Q_4$ | 3312056 | 21237 | 15436 | 0.9954 | 0.9936 | 0.9945 |
| | $Q_2$ | 3310979 | 21442 | 16513 | 0.9950 | 0.9936 | 0.9943 |
| | $F_{10}$ | 3308694 | 21260 | 18798 | 0.9944 | 0.9936 | 0.9940 |
| | $\langle Q_2, Q_8 \rangle_\delta$ | 3311573 | 21820 | 15919 | 0.9952 | 0.9935 | 0.9943 |
| 30× | Original | 3307863 | 60007 | 19629 | 0.9941 | 0.9822 | 0.9881 |
| | $Q_8$ | 3307922 | 63022 | 19570 | 0.9941 | 0.9813 | 0.9877 |
| | $Q_4$ | 3307467 | 62382 | 20025 | 0.9940 | 0.9815 | 0.9877 |
| | $Q_2$ | 3306457 | 61682 | 21035 | 0.9937 | 0.9817 | 0.9876 |
| | $F_{10}$ | 3303993 | 61874 | 23499 | 0.9929 | 0.9816 | 0.9872 |
| | $\langle Q_2, Q_8 \rangle_\delta$ | 3306928 | 62256 | 20564 | 0.9938 | 0.9815 | 0.9876 |
| 20× | Original | 3285668 | 466868 | 41824 | 0.9874 | 0.8756 | 0.9282 |
| | $Q_8$ | 3285516 | 494441 | 41976 | 0.9874 | 0.8692 | 0.9245 |
| | $Q_4$ | 3284787 | 464226 | 42705 | 0.9872 | 0.8762 | 0.9284 |
| | $Q_2$ | 3283171 | 432505 | 44321 | 0.9867 | 0.8836 | 0.9323 |
| | $F_{10}$ | 3277795 | 442196 | 49697 | 0.9851 | 0.8812 | 0.9302 |
| | $\langle Q_2, Q_8 \rangle_\delta$ | 3283622 | 442501 | 43870 | 0.9868 | 0.8813 | 0.9311 |

**Table 7:** Metrics TP, FP, FN, Recall, Precision, and F1 for the INDELs obtained with PEPPER-Margin-DeepVariant under various coverage scenarios, for the original (non-quantized) data and for the data obtained by applying various quality score quantizers.

| Coverage | Quantization | TP | FP | FN | Recall | Precision | F1 |
| --- | --- | --- | --- | --- | --- | --- | --- |
| 90× | Original | 303517 | 29403 | 200983 | 0.6016 | 0.9136 | 0.7255 |
| | $Q_8$ | 303713 | 29607 | 200787 | 0.6020 | 0.9131 | 0.7256 |
| | $Q_4$ | 301600 | 30180 | 202900 | 0.5978 | 0.9110 | 0.7219 |
| | $Q_2$ | 296238 | 30968 | 208262 | 0.5872 | 0.9074 | 0.7130 |
| | $F_{10}$ | 289838 | 28569 | 214662 | 0.5745 | 0.9123 | 0.7050 |
| | $\langle Q_2, Q_8 \rangle_\delta$ | 304333 | 37142 | 200167 | 0.6032 | 0.8935 | 0.7202 |
| 50× | Original | 287895 | 43665 | 216605 | 0.5707 | 0.8709 | 0.6895 |
| | $Q_8$ | 287890 | 44025 | 216610 | 0.5706 | 0.8699 | 0.6892 |
| | $Q_4$ | 286319 | 43663 | 218181 | 0.5675 | 0.8703 | 0.6870 |
| | $Q_2$ | 283147 | 44307 | 221353 | 0.5612 | 0.8673 | 0.6815 |
| | $F_{10}$ | 279418 | 42311 | 225082 | 0.5539 | 0.8711 | 0.6772 |
| | $\langle Q_2, Q_8 \rangle_\delta$ | 289053 | 51537 | 215447 | 0.5729 | 0.8515 | 0.6850 |
| 40× | Original | 278841 | 51239 | 225659 | 0.5527 | 0.8476 | 0.6691 |
| | $Q_8$ | 278920 | 51314 | 225580 | 0.5529 | 0.8475 | 0.6692 |
| | $Q_4$ | 277724 | 51171 | 226776 | 0.5505 | 0.8473 | 0.6674 |
| | $Q_2$ | 275492 | 51998 | 229008 | 0.5461 | 0.8442 | 0.6632 |
| | $F_{10}$ | 272761 | 50237 | 231739 | 0.5407 | 0.8474 | 0.6601 |
| | $\langle Q_2, Q_8 \rangle_\delta$ | 280046 | 59427 | 224454 | 0.5551 | 0.8281 | 0.6646 |
| 30× | Original | 265283 | 63876 | 239217 | 0.5258 | 0.8092 | 0.6374 |
| | $Q_8$ | 265193 | 64151 | 239307 | 0.5257 | 0.8085 | 0.6371 |
| | $Q_4$ | 264442 | 64213 | 240058 | 0.5242 | 0.8079 | 0.6358 |
| | $Q_2$ | 263047 | 65037 | 241453 | 0.5214 | 0.8051 | 0.6329 |
| | $F_{10}$ | 260681 | 63727 | 243819 | 0.5167 | 0.8069 | 0.6300 |
| | $\langle Q_2, Q_8 \rangle_\delta$ | 266429 | 72561 | 238071 | 0.5281 | 0.7894 | 0.6329 |
| 20× | Original | 239522 | 96040 | 264978 | 0.4748 | 0.7177 | 0.5715 |
| | $Q_8$ | 239532 | 96766 | 264968 | 0.4748 | 0.7162 | 0.5710 |
| | $Q_4$ | 238938 | 95403 | 265562 | 0.4736 | 0.7185 | 0.5709 |
| | $Q_2$ | 237475 | 94862 | 267025 | 0.4707 | 0.7185 | 0.5688 |
| | $F_{10}$ | 235169 | 92383 | 269332 | 0.4661 | 0.7219 | 0.5665 |
| | $\langle Q_2, Q_8 \rangle_\delta$ | 239968 | 102876 | 264532 | 0.4757 | 0.7039 | 0.5677 |

**Figure 1:**
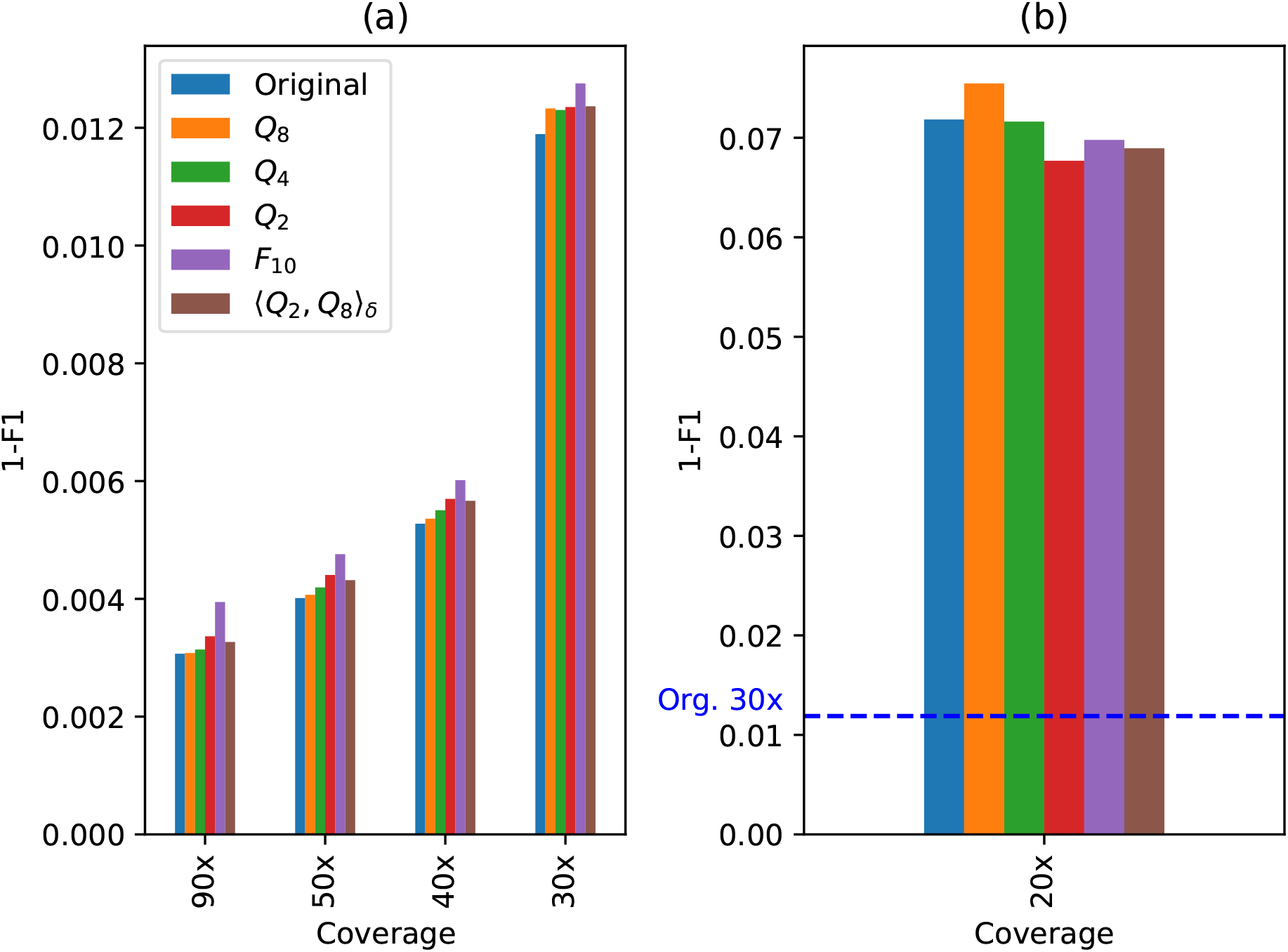
Plot of 1 – F1 score for the SNPs obtained with PEPPER-Margin-DeepVariant for the original (non-quantized) data and for the data obtained by applying various quality score quantizers, for coverages 90×, 50×, 40×, and 30× in plot (a), and for 20× in plot (b). As a reference, the dotted line in (b) marks the value 1 – F1 for the original data with coverage 30×.

**Figure 2:**
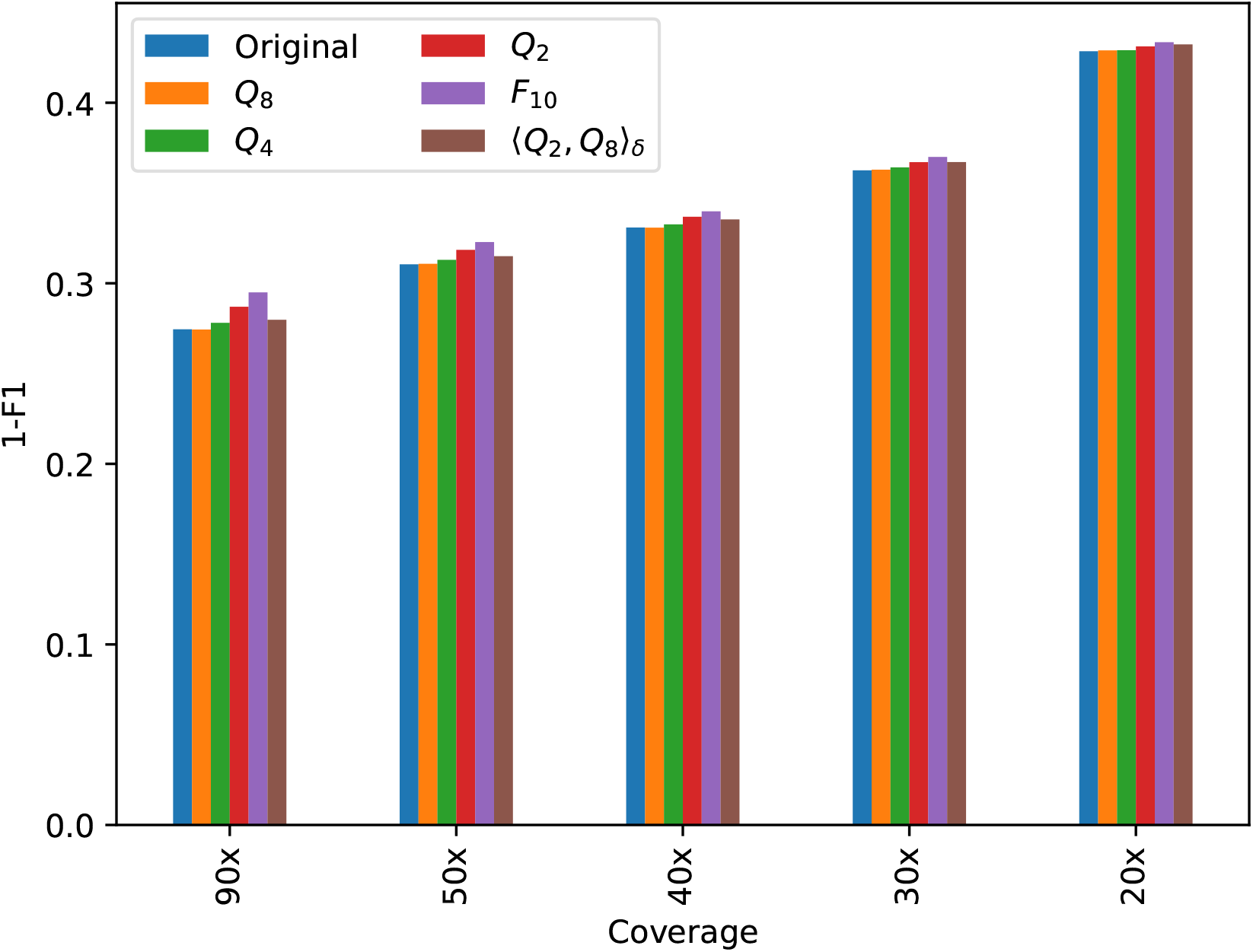
Plot of 1 – F1 score for the INDELs obtained with PEPPER-Margin-DeepVariant under various coverage scenarios, for the original (non-quantized) data and for the data obtained by applying various quality score quantizers.

For SNPs, Table 6 shows high recall and precision values, specially for large coverage levels. In all cases, the difference b etween t he F1 score obtained with the original data and that obtained with a quantized version of the same data is rather small. The largest difference f or c overage 3 0× a nd a bove is smaller than 10^−3^ (for *F*_10_ on 90× data). For each of these coverage leveles, the performance slightly improves in general with the quantization precision, and the performance for ⟨*Q*_2_, *Q*_8_⟩_*δ*_ lies between that of *Q*_2_ and *Q*_4_ (see Figure 1 (a)). For the shallowest coverage, 20×, the performance is noticeable worse (see Figure 1 (b) and note the different scales in plots (a) and (b) of the figure). In this case the F1 score not always improves with quantization precision. Following the performance comparison criteria in [13], we notice that PEPPER-Margin-DeepVariant on 90× data still outperforms DeepVariant on Illumina short reads (35× Illumina NovaSeq, F1: 0.9963) for all quantizers except *F*_10_. Another interesting reference for comparison is the variation in performance obtained by using different v ariant c alling p ipelines. For e xample, using Longshot [22] on the 30× original data we obtain a precision of 0.9648 and a recall of 0.9743, which determine an F1 score of 0.9695 for SNPs. Since Longshot ignores quality scores, the results after quality score quantization are exactly the same. Notice that these metric values fall below those reported in Table 6 for every quantizer on 30× data, including *F*_10_, which losses all quality score information from the original data. Thus, in this example, the choice of a variant calling pipeline determines larger performance variations than the quantization of quality scores.

For INDELs, in agreement with [13], the recall and precision values are significantly worse than those obtained for SNPs. The difference between the performance obtained with non-quantized and quantized data is still small, although more noticeable than the difference obtaind for SNPs. For INDELs, the largest difference in F 1 score occurs for *F* _10_ on 9 0× data, with a difference approximately equal to 0. 002. For *Q*_4_, the maximum difference drops to 3.6 × 10^−3^, and for *Q*_8_ it is 4.7 × 10^−4^. The variant calling performance for the data quantized with ⟨*Q*_2_, *Q*_8_⟩_*δ*_ lies, in general, between those obtained with *Q*_2_ and *Q*_4_.

Table 8 shows the compression ratio obtained with gzip for various quantizers. The table also shows the space saving obtained by quality score quantization, calculated as the difference in size between the compressed original data set and the compressed quantized data set, expressed as a percent with respect to the size of the compressed original data set. For example, the table shows that gzip compresses the original data set to roughly half the original size (the compression ratio is 0.49), and *Q*_4_ improves the compression ratio to 0.25, yielding a compressed data set that is roughly half the size of the compressed original data set (a saving of 48.52 %). For the most heavy quantizer, *F*_10_, the difference in compressed size represents almost 70% of the original compressed data set, which in the evaluated data set amounts to more than 175 GB of saved space. For easy assessment of the relation between compression ratio and variant calling performance, Table 8 shows, for each quantizer, the F1 score obtained with quantized 90× coverage data.

**Table 8:** Compression ratio obtained with gzip for the original data and for various quantized versions of the same data. The third column presents the percent space saving of the gzip compression of each quantized version compared to the original data compressed with gzip. The right-most column shows, for each quantization level, the F1 score obtained with coverage level 90×.

| Quantization | Compression ratio | Quantization space saving (%) | F1 score at $90\times$ |
| --- | --- | --- | --- |
| Original | 0.49 | — | 0.9969 |
| $Q_8$ | 0.32 | 33.73 | 0.9969 |
| $Q_4$ | 0.25 | 48.52 | 0.9969 |
| $Q_2$ | 0.19 | 60.70 | 0.9966 |
| $F_{10}$ | 0.15 | 69.30 | 0.9961 |
| $\langle Q_2, Q_8 \rangle_\delta$ | 0.22 | 54.89 | 0.9967 |

The trade off between storage space and variant calling performance is also illustrated in Figure 3. For each quantizer and coverage level *ℓ* (excluding 20×), we multiply the gzip compression ratio for that quantizer by the coverage level ratio, *ℓ*/90. This value represents the fraction of storage space required by the data with coverage level *ℓ*, quantized and compressed with gzip, with respect to the full (uncompressed, 90×) data set. These fractions of storage space are plotted in the figure against the F1 score obtained for SNPs in each case. Notice, for example, that higher coverage data quantized with *Q*_4_ yields better performance than lower coverage non-quantized data (except at 90× coverage), requiring actually less storage space.

**Figure 3:**
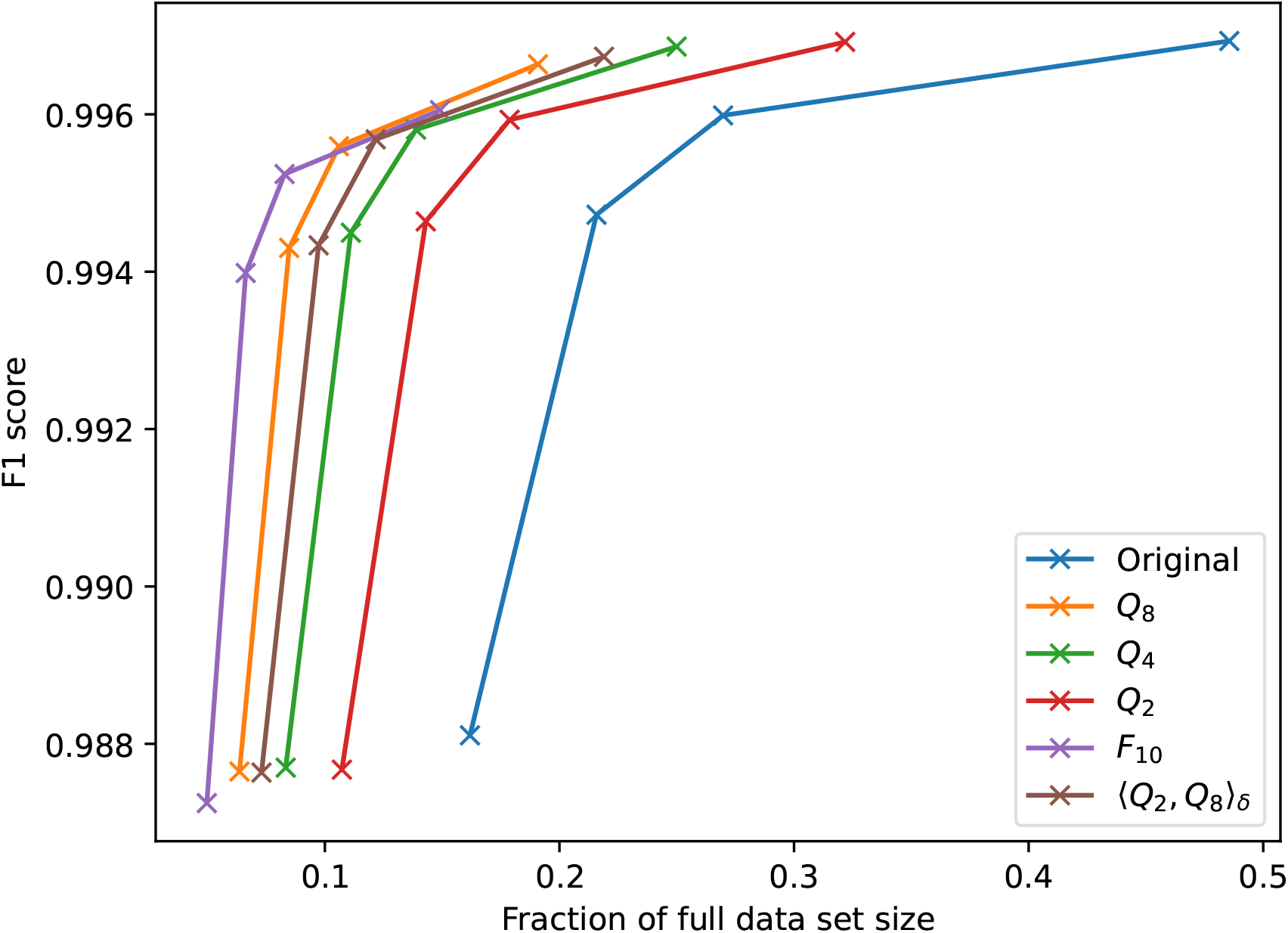
F1 score for the SNPs obtained with PEPPER-Margin-DeepVariant as a function of the required storage space, for each evaluated quantizer and coverage levels 90×, 50×, 40×, and 30×. The storage requirement is expressed as a fraction of the size of the full (uncompressed, 90×) data set.

## 4 Conclusions

Our experiments on various usage scenarios for nanopore sequencing data, including different applications and coverage levels, show that the precision that is currently used for quality scores is unnecessarily high. A quantizer like *Q*_4_, wich uses only four different values for quality scores, yields results similar to those obtained with the original data in all tested scenarios. We point out that all these results were obtained with applications *as they are provided*, with no special tuning or training for quantized quality scores. Although such specific tuning may improve the performance of these applications (for example through neural network retraining), the matter of fact is that excelent results are obtained with no software adjustment. The quantization of quality scores results in large storage space savings, even using a general purpose compressor such as gzip.

## 5 Acknowledgments

This work has been partially founded by ANII, grant FSDA 1 2018 1 154790. Some of the experiments presented in this paper were carried out using ClusterUY (site: https://cluster.uy).

## Footnotes

1 https://nanoporetech.com/

2 https://github.com/nanoporetech/medaka

3 https://nanopore.s3.climb.ac.uk/mock/Zymo-GridION-EVEN-3Peaks-R103-merged.fq.gz

4 https://github.com/nanoporetech/medaka

5 http://nanopore.s3.climb.ac.uk/mockcommunity/v2/Zymo-Isolates-SPAdes-Illumina.fasta

6 http://github.com/illumina/hap.py

7 https://www.gnu.org/software/gzip/

## References

[1] “Novaseq™ 6000 system quality scores and RTA3 software,” technical report, Illumina, 2017.

[2] “Reducing whole-genome data storage footprint,” technical report, Illumina, 2014.

[3] I. Ochoa, H. Asnani, D. Bharadia, M. Chowdhury, T. Weissman, and G. Yona, “Qualcomp: a new lossy compressor for quality scores based on rate distortion theory,” BMC Bioinformatics, vol. 14, no. 1, 2013.

[4] R. Cánovas, A. Moffat, and A. Turpin, “Lossy compression of quality scores in genomic data,” Bioinformatics, vol. 30, no. 15, pp. 2130–2136, 2014.

[5] I. Ochoa, M. Hernaez, R. Goldfeder, T. Weissman, and E. Ashley, “Effect of lossy compression of quality scores on variant calling,” Briefings in bioinformatics, vol. 18, no. 2, pp. 183–194, 2016.

[6] G. Dufort y Álvarez, G. Seroussi, P. Smircich, J. Sotelo, I. Ochoa, and Á. Martín, “ENANO: Encoder for NANOpore FASTQ files,” Bioinformatics, vol. 36, pp. 4506–4507, 05 2020.

[7] G. Dufort y Álvarez, G. Seroussi, P. Smircich, J. Sotelo-Silveira, I. Ochoa, and Á. Martín, “RENANO: a REference-based compressor for NANOpore FASTQ files,” Bioinformatics, 06 2021.

[8] Q. Meng, S. Chandak, Y. Zhu, and T. Weissman, “Nanospring: reference-free lossless compression of nanopore sequencing reads using an approximate assembly approach,” bioRxiv, 2021.

[9] M. Kokot, A. Gudyś, H. Li, and S. Deorowicz, “CoLoRd: Compressing long reads,” bioRxiv, 2021.

[10] R. Vaser, I. Sović, N. Nagarajan, and M. Šikić, “Fast and accurate de novo genome assembly from long uncorrected reads,” Genome Res, vol. 27, pp. 737–746, 05 2017.

[11] R. Poplin, P.-C. Chang, D. Alexander, S. Schwartz, T. Colthurst, A. Ku, D. Newburger, J. Dijamco, N. Nguyen, P. T. Afshar, S. S. Gross, L. Dorfman, C. Y. McLean, and M. A. DePristo, “A universal snp and small-indel variant caller using deep neural networks,” Nature Biotechnology, vol. 36, pp. 983–987, Nov 2018.

[12] K. Shafin, T. Pesout, R. Lorig-Roach, M. Haukness, H. E. Olsen, C. Bosworth, J. Armstrong, K. Tigyi, N. Maurer, S. Koren, F. J. Sedlazeck, T. Marschall, S. Mayes, V. Costa, J. M. Zook, K. J. Liu, D. Kilburn, M. Sorensen, K. M. Munson, M. R. Vollger, J. Monlong, E. Garrison, E. E. Eichler, S. Salama, D. Haussler, R. E. Green, M. Akeson, A. Phillippy, K. H. Miga, P. Carnevali, M. Jain, and B. Paten, “Nanopore sequencing and the shasta toolkit enable efficient de novo assembly of eleven human genomes,” Nature Biotechnology, vol. 38, pp. 1044–1053, Sep 2020.

[13] K. Shafin, T. Pesout, P.-C. Chang, M. Nattestad, A. Kolesnikov, S. Goel, G. Baid, M. Kolmogorov, J. M. Eizenga, K. H. Miga, P. Carnevali, M. Jain, A. Carroll, and B. Paten, “Haplotype-aware variant calling with pepper-margin-deepvariant enables high accuracy in nanopore long-reads,” Nature Methods, vol. 18, pp. 1322–1332, Nov 2021.

[14] C. Delahaye and J. Nicolas, “Sequencing DNA with nanopores: Troubles and biases,” PLOS ONE, vol. 16, pp. 1–29, 10 2021.

[15] S. M. Nicholls, J. C. Quick, S. Tang, and N. J. Loman, “Ultra-deep, long-read nanopore sequencing of mock microbial community standards,” GigaScience, vol. 8, 05 2019.

[16] M. Kolmogorov, D. M. Bickhart, B. Behsaz, A. Gurevich, M. Rayko, S. B. Shin, K. Kuhn, J. Yuan, E. Polevikov, T. P. L. Smith, and P. A. Pevzner, “metaflye: scalable long-read metagenome assembly using repeat graphs,” Nature Methods, vol. 17, pp. 1103–1110, Nov 2020.

[17] A. Bankevich, S. Nurk, D. Antipov, A. A. Gurevich, M. Dvorkin, A. S. Kulikov, V. M. Lesin, S. I. Nikolenko, S. Pham, A. D. Prjibelski, A. V. Pyshkin, A. V. Sirotkin, N. Vyahhi, G. Tesler, M. A. Alekseyev, and P. A. Pevzner, “SPAdes: a new genome assembly algorithm and its applications to single-cell sequencing,” J Comput Biol, vol. 19, pp. 455–477, May 2012.

[18] A. Mikheenko, V. Saveliev, and A. Gurevich, “MetaQUAST: evaluation of metagenome assemblies,” Bioinformatics, vol. 32, pp. 1088–1090, 11 2015.

[19] J. Ruan and H. Li, “Fast and accurate long-read assembly with wtdbg2,” Nature Methods, vol. 17, pp. 155–158, Feb 2020.

[20] G. P. Consortium, A. Auton, L. D. Brooks, R. M. Durbin, E. P. Garrison, H. M. Kang, J. O. Korbel, J. L. Marchini, S. McCarthy, G. A. McVean, and G. R. Abecasis, “A global reference for human genetic variation,” Nature, vol. 526, pp. 68–74, Oct 2015.

[21] A. Gurevich, V. Saveliev, N. Vyahhi, and G. Tesler, “QUAST: quality assessment tool for genome assemblies,” Bioinformatics, vol. 29, pp. 1072–1075, 02 2013.

[22] P. Edge and V. Bansal, “Longshot enables accurate variant calling in diploid genomes from single-molecule long read sequencing,” Nature Communications, vol. 10, p. 4660, Oct 2019.

